# The timing of confidence computations in human prefrontal cortex

**DOI:** 10.1101/2023.03.21.533662

**Authors:** Kai Xue, Yunxuan Zheng, Farshad Rafiei, Dobromir Rahnev

**Affiliations:** School of Psychology, Georgia Institute of Technology, Atlanta, GA

**Keywords:** confidence, DLPFC, metacognition, perceptual decision making, TMS

## Abstract

Knowing when confidence computations take place is critical for building mechanistic understanding of the neural and computational bases of metacognition. Yet, even though substantial amount of research has focused on revealing the neural correlates and computations underlying human confidence judgments, very little is known about the timing of confidence computations. Subjects judged the orientation of a briefly presented visual stimulus and provided a confidence rating regarding the accuracy of their decision. We delivered single pulses of transcranial magnetic stimulation (TMS) at different times after stimulus presentation. TMS was delivered to either dorsolateral prefrontal cortex (DLPFC) in the experimental group or to vertex in the control group. We found that TMS to DLPFC, but not to vertex, led to increased confidence in the absence of changes to accuracy or metacognitive ability. Critically, equivalent levels of confidence increase occurred for TMS delivered between 200 and 500 ms after stimulus presentation. These results suggest that confidence computations occur during a broad window that begins before the perceptual decision has been fully made and thus provide important constraints for theories of confidence generation.

## Introduction

Metacognition, the ability to assess the quality of our own decisions, is crucial for effective decision-making (Fleming et al., 2012; Koriat, 2007, 2007; Metcalfe & Shimamura, 1994; Nelson, 1990; Shimamura, 2000; Yeung & Summerfield, 2012). Substantial amount of research has focused on revealing the neural correlates underlying human confidence judgments (AbdulSabur et al., 2014; Fleming et al., 2012; Morales et al., 2018; Pereira et al., 2017; Shekhar & Rahnev, 2018; Shimamura, 2000; Yeon et al., 2020; Zheng et al., 2021). Many studies have pointed to a central role for the prefrontal cortex (Janowsky et al., 1989; Shimamura, 2000; Shimamura & Squire, 1986) and specifically the dorsolateral prefrontal cortex (DLPFC), which has been linked to confidence judgements (Fleming et al., 2012; Rounis et al., 2010; Shekhar & Rahnev, 2018).

However, although much progress has been made in discovering where confidence is computed in the brain, much less is known about the timing of confidence computation (Desender, Donner, et al., 2021; Dotan et al., 2018; Fetsch et al., 2014, 2018; Moran et al., 2015; Pleskac & Busemeyer, 2010). Recently, Shekhar and Rahnev (2018) examined this question by delivering a train of three pulses of transcranial magnetic stimulation (TMS) to DLPFC at 250, 350, and 450 ms after stimulus onset in a perceptual decision-making task. They found that the TMS train of pulses decreased confidence compared to a control region (the primary somatosensory cortex) but could not determine exactly when confidence computation occurred besides the fact that some part of the window between 250 and 450 ms after stimulus onset is important.

To address more precisely the issue of when confidence computations occur, here we used single pulse TMS at four different times. Specifically, we delivered single pulses of TMS at 200, 300, 400, and 500 ms after stimulus onset and compared the results to TMS delivered simultaneously with stimulus onset (0 ms condition). Subjects judged the orientation of a briefly presented visual stimulus and reported their confidence. We delivered online TMS to DLPFC in the experimental group, and to vertex in the control group. We found that TMS to DLPFC, but not to vertex, led to an increase in confidence without any changes to accuracy or metacognitive ability. More importantly, the levels of confidence increase brought by TMS were the same across intervals between 200 and 500 ms after the stimulus presentation. These results suggest that confidence computations occur during a broad time window. Because the perceptual decision is unlikely to be made within 200 ms on a substantial proportion of trials, these results go against strong versions of the post-decisional theories of confidence where all confidence computations happen only after the decision has already been made.

## Methods

### Preregistration

We preregistered the sample size, exclusion criteria, and analyses for the DLPFC TMS group (https://osf.io/3ru2m). After the data for the DLPFC group were collected, we additionally collected data from a control group where we targeted vertex instead of DLPFC.

### Subjects

A total of 76 subjects were enrolled in the study with 50 subjects in the experiment group (TMS to DLPFC) and 26 subjects in the control group (TMS to vertex). Based on our preregistered criteria, we excluded a total of 13 subjects. Specifically, we excluded nine subjects who did not finish the experiment either because of TMS-related discomfort (six subjects) or because they did not complete all trials before the end of the session (three subjects). Another subject was excluded because their data was lost because of computer malfunction. Finally, we excluded three subjects for performance lower than 55% correct. Thus, the final sample size consisted of 62 subjects (22 females and 40 males) with 43 subjects in the experimental and 19 subjects in the control group. All subjects were right-handed, had normal or corrected-to-normal vision, and had no history of seizure, family history of epilepsy, stroke, severe headache, or metal anywhere in the head. All subjects provided informed consent and were compensated $30 for two hours of total participation.

### Task

Each trial began with subjects fixating on a small white dot (size = 0.05°) at the center of the screen for 500 ms, followed by presentation of a Gabor patch (diameter = 3°) oriented either to the right (clockwise, 45°) or to the left (counterclockwise, 135°) of vertical for 100 ms. The Gabor patch was superimposed on a noisy background. Subjects indicated the orientation of the Gabor patch while simultaneously rating their confidence on a 4-point scale (where 1 corresponds to lowest confidence and 4 corresponds to highest confidence) via a single key press (Figure 1A). The four fingers of the left hand were mapped to the four confidence ratings for the left tilt response, whereas the four fingers of the right hand were mapped to the four confidence ratings for the right tilt response. For each hand, the index finger indicated a confidence of 1, whereas the pinky finger indicated a confidence of 4. The orientation of the stimulus (left/right) was chosen randomly on each trial.

**Figure 1.**
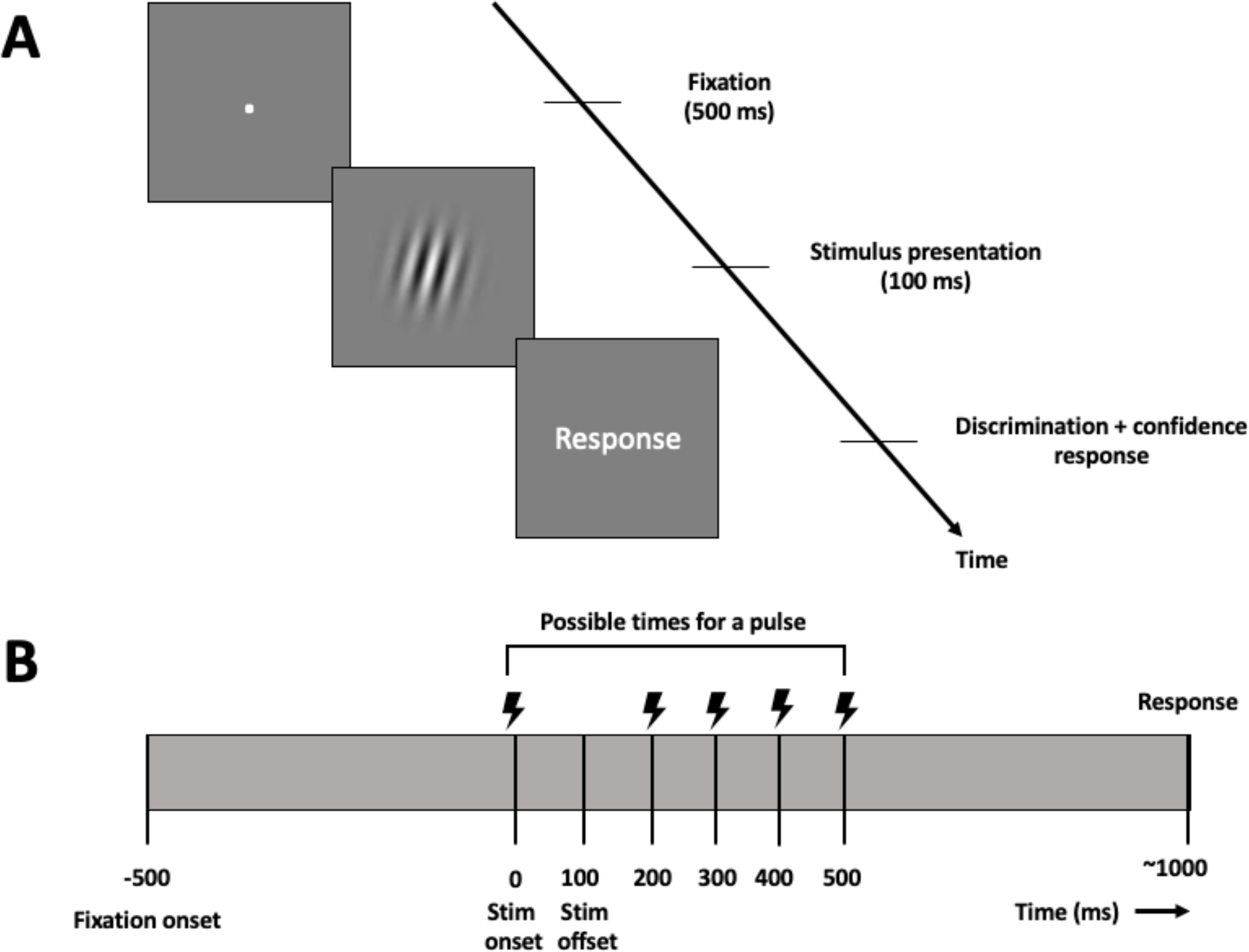
Task. (A) Trial sequence. Each trial began with a short fixation (500 ms), followed by the presentation of an oriented Gabor patch (100 ms). Subjects had to simultaneously indicate the tilt (left/right) of the Gabor patch and their confidence on a 4-point scale. (B) Timeline of TMS delivery. TMS was delivered as a single pulse 0, 200, 300, 400, or 500 ms after the stimulus onset. Subjects had a mean response time of 1078 ms.

We delivered online TMS as a single pulse on each trial at 0, 200, 300, 400, or 500 ms after the stimulus onset. We chose the 200-500 ms delays to coincide with the presumed time window of confidence computation. Indeed, in a previous study, we delivered TMS to DLPFC as a train of three pulses starting at 250 and ending at 450 ms after the stimulus onset, and found an effect on confidence (Shekhar & Rahnev, 2018). Further, neuronal recordings from monkeys suggest that the discrimination response emerges about 200 ms after stimulus onset (Siegel et al., 2015), suggesting that confidence computation in human DLPFC are unlikely to happen much earlier than 200 ms. The 0-ms condition was chosen as a control against which to compare the four delay conditions.

### Design and procedure

The main experiment consisted of four runs each consisting of five 40-trial blocks (for a total of 800 trials). The five possible TMS delays (0, 200, 300, 400, and 500 ms after the stimulus onset) were presented in a pseudorandom order such that within each group of five trials, each delay appeared once. The design and procedure were identical for the DLPFC and vertex groups except for the targeted site.

At the beginning of the experiment, subjects underwent a behavioral training procedure without TMS. The training session started with high Gabor patch contrast value (80%) and gradually progressed to lower contrast values (the last block had contrast values of 6%). Subjects were given trial-by-trial feedback on their performance during this training period. Then, subjects completed a 3-down-1-up staircasing procedure without feedback to adaptively estimate the contrast for each individual subject (Macmillan & Creelman, 2004). This procedure yielded a mean contrast value of 6.64% (SD = 0.96%). We used the contrast value obtained for each subject for the rest of the experiment.

At the end of the training, subjects completed one practice TMS block with the same level of contrast and TMS delivery as in the rest of the experiment. The practice block was included to accustom subjects to receiving TMS while performing the task. The practice block was excluded from further analyses.

### Apparatus

The stimuli were generated using Psychophysics Toolbox (RRID: SCR_002881) in MATLAB (The MathWorks, RRID:SCR_001622). During the training and the main experiment, subjects were seated in a dim room and were positioned 60 cm away from the computer screen (21.5-inch display, 1920 × 1080 pixel resolution, 60 Hz refresh rate).

### Defining the targets for TMS targeting

We defined two sites as targets for TMS: DLPFC and vertex. Based on previous studies, we localized right DLPFC using the location of the F4 electrode in the 10-20 system used for EEG electrode placement (Conson et al., 2015; Fitzgerald, 2021; Fitzgerald et al., 2009; Mir-Moghtadaei et al., 2015; Rusjan et al., 2010). As in previous studies that targeted DLPFC with TMS during perceptual decision-making tasks (Rahnev et al., 2016; Shekhar & Rahnev, 2018), the DLPFC target was defined in the right hemisphere because the right hemisphere is dominant for visual processing (Hellige, 1996).

To determine the subject-specific location for stimulation, we followed the Beam F3 Location System developed by Beam and Borckardt (2009). This method allowed us to precisely determine the F4 region using skull measurements (Beam et al., 2009). The location of the vertex was determined as the midpoint between the Nasion and inion.

### TMS setup

TMS was delivered with a magnetic stimulator (MagPro R100; MagVenture, RRID:SCR_009601) using a figure-eight coil with a diameter of 75 mm. We determined the resting motor threshold (RMT) immediately before starting the main experiment. To localize the motor cortex, we marked its putative location and applied suprathreshold single pulses around that location. We determined the location of the right motor cortex as the region that induced maximal twitches of the fingers in the left hand. Then, using this location as the target, we determined the RMT using an adaptive parameter estimation by sequential testing procedure (Borckardt et al., 2006). For one subject, we were unable to estimate RMT reliably, so this subject was excluded from the experiment.

The TMS coil was oriented tangential to the skull such that the induced magnetic field was orthogonal to the skull. Stimulation was delivered at 120% of the individual RMT (average stimulation intensity = 72% of maximum stimulator output). In two cases, the stimulation intensity exceeded 80% of maximum stimulator output. Due to discomfort, the intensity was reduced to 80% of maximum stimulator output for both subjects. No arm or leg movements were elicited by stimulation of either DLPFC or vertex.

### Analyses

We analyzed the accuracy, reaction time (RT), confidence, and metacognitive ability for each delay condition. The metacognitive ability was operationalized using the measure Mratio developed by Maniscalco and Lau (2012). Mratio is derived from signal detection theoretical modeling of the observer’s decision and confidence responses. It is the ratio of two measures: the observer’s metacognitive sensitivity (*meta-d’*, the ability to discriminate between correct and incorrect responses) and the observer’s stimulus sensitivity (*d’*, the ability to discriminate between the two stimulus classes). The ratio of meta-d’ to d’ factors out the contribution of stimulus sensitivity towards metacognitive performance and captures the efficiency of the observer’s metacognitive processes (Fleming & Lau, 2014).

To examine the effect of TMS on confidence and metacognitive ability (Mratio), we computed the difference between the confidence and the Mratio scores of each delay condition and 0-ms condition. Then, we compared the obtained difference scores between two TMS stimulation sites (DLPFC and vertex) and the four TMS delay conditions (200, 300, 400, and 500 ms) using one-way and two-way repeated-measures ANOVAs. Direct comparisons between the two TMS stimulation sites within each delay condition were made using independent sample t-tests, whereas direct comparisons between different delay conditions within a single stimulation site were made using paired t-tests.

Note that the analyses performed above differ in some ways from the analyses we preregistered. The reason for this is that our preregistration anticipated that there would be differences between the TMS effects for the four different delay conditions, and that there may be individual variability between subjects as to the most effective delay condition. Because neither of these assumptions was supported by the data, the analyses we preregistered are subsumed within the simpler analyses we performed instead. Nevertheless, for completeness, we report the results of all preregistered analyses in the Supplementary Results.

### Data and code

All data and code are available at https://osf.io/szr9u/.

## Results

We investigated the timing of confidence computation in DLPFC. To do so, we used an online TMS protocol to disrupt DLPFC activity at different time points after stimulus onset and compared the effects to a control condition where TMS was delivered over vertex. Subjects indicated the tilt (left/right) of a noisy Gabor patch while simultaneously providing confidence rating on a 4-point scale. On each trial, we delivered a single pulse of TMS at one of the four possible delays (200, 300, 400, and 500 ms) after the stimulus onset and compared the results to a condition where TMS was delivered at stimulus onset (i.e., 0 ms delay).

Previous work consistently found that TMS to DLPFC had no effect on either accuracy or reaction time (RT) (Rahnev et al., 2016; Rounis et al., 2010; Ryals et al., 2016; Shekhar & Rahnev, 2018). However, we observed that the 0-ms condition in the DLPFC group produced lower accuracy (74.1% correct) than the four delay conditions (200 ms: 76.5% correct, t(42) = 3.42, *p* = 0.001; 300 ms: 75.5% correct, t(42) = 1.68, *p* = 0.04; 400 ms: 76.2% correct, t(42) = 3.09, *p* = 0.002; 500 ms: 76.1% correct, t(42) = 2.59, *p* = 0.007). These results appear consistent with the notion that the 0 ms TMS may have induced an eye blink or otherwise interfered with the initial processing of the stimulus. Because the decrease in accuracy for the 0-ms condition occurred for some subjects only, for subsequent analyses we excluded all subjects for whom the 0-ms condition had accuracy more than 3.5% lower than the average of the four delay conditions. This led to the exclusion of 12 subjects in the DLPFC group while also equating the accuracy of the 0-ms condition (average accuracy = 75.69%) and average accuracy in the four delay conditions (average accuracy = 75.76%; t(30) = 0.18, *p* = 0.86). We also applied the same exclusion criterion to the vertex group, which led to the exclusion of 3 subjects and also equated the average accuracy of the 0-ms condition (average accuracy = 75.98%) and the average accuracy in the four delay conditions (average accuracy = 76.11%; t(15) = -1.42, *p* = 0.17). The lower rate of exclusion for the vertex condition is consistent with the possibility that TMS at 0 ms may have induced eye blinks for some subjects, but this happened primarily for DLPFC since that site is closer to the eye sockets. Repeating the analyses below without these exclusions still leads to the same main conclusions (Supplementary Figures 1-3).

We then examined the effects of TMS on task performance across the four delay conditions. A two-way repeated-measures ANOVA on the accuracy difference between each delay condition and the 0-ms baseline condition with factors TMS site (DLPFC and vertex) and delay conditions (200, 300, 400, and 500 ms) showed that there was no main effect of TMS site (F_(1,180)_ = 1.91, *p* = 0.13), no main effect of delay condition (F_(3,180)_ = 1.3, *p* = 0.13), and no interaction between TMS site and delay condition (F_(3,180)_ = 0.39, *p* = 0.76; Figure 2A). A similar two-way repeated-measures ANOVA on the RT difference between each delay condition and the 0-ms baseline condition also showed no effect of TMS site (F_(1,180)_ = 2.0, *p* = 0.16), delay condition (F_(3,180)_ = 0.24, *p* = 0.87), or an interaction between the two (F_(3,180)_ = 0.90, *p* = 0.90; Figure 2B). Pairwise comparisons between the DLPFC TMS and vertex TMS groups for each delay condition also showed no differences in either accuracy or RT (all *p*’s > 0.14). Thus, TMS had equivalent effects at delays between 200 and 500 ms for both accuracy and RT.

**Figure 2.**
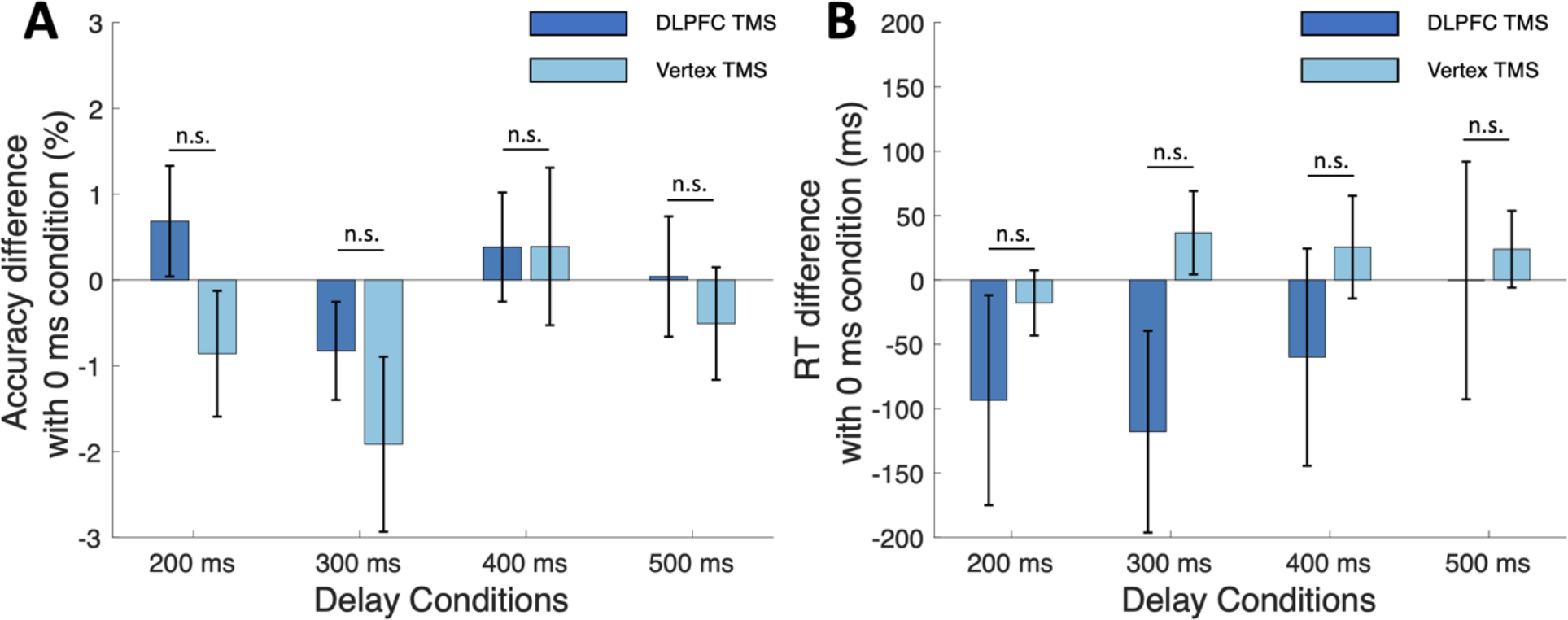
TMS effects on accuracy and RT. (A) The effect of TMS on the accuracy difference between each delay condition and the 0-ms condition. The accuracy difference does not depend on either the TMS site, the delay condition, or the interaction between TMS site and the delay condition. (B) The effect of TMS on RT difference between each delay condition and 0-ms condition. The RT difference does not depend on either the TMS site, the delay condition, or the interaction between TMS site and the delay condition. Error bars represent SEM; n.s., *p* > .05.

Having established that the four delay conditions do not differentially affect performance, we examined whether TMS with different timing had differential effect on confidence or metacognitive efficiency. We originally hypothesized that DLPFC TMS will lead to decrease in confidence and that this effect will be stronger in some delay conditions than others. However, the results showed opposite patterns for both of these hypotheses. First, instead of a decrease, TMS to DLPFC led to an increase in confidence for each delay condition compared to the 0-ms condition (200 ms: t(30) = 5.37, *p* = 4.15 × 10^−6^; 300 ms: t(30) = 5.23, *p* = 5.99 × 10^−6^; 400 ms: t(30) = 5.60, *p* = 2.14 × 10^−6^; 500 ms: t(30) = 4.75, *p* = 2.38 × 10^−5^; Figure 3). This effect was not present when TMS was delivered to the vertex (all *p*’s > 0.86). Further, all pairwise comparisons between DLPFC and vertex TMS showed significant differences in confidence (*p* < 0.001 for all four comparisons). Second, instead of the hypothesized differences among the four delay conditions, we found that the increase in confidence was equivalent for all four conditions. Indeed, a one-way ANOVA on the confidence in the 200-500 ms delay conditions for the DLPFC TMS group showed no significant effect of condition (F_(3,120)_ = 0.03, *p* = 0.99). A similar one-way ANOVA for the vertex group also showed no significant effect of condition (F_(3,60)_ = 0.28, *p* = 0.83). Direct comparisons between any pair of delay conditions for both the DLPFC TMS and vertex TMS groups confirmed the lack of any significant differences between the delay conditions (*p* > 0.09 for all 12 pairwise comparisons). These results confirmed previous findings suggesting that DLPFC is involved in confidence computation (Shekhar & Rahnev, 2018), and further demonstrate that the computation doesn’t happen within a narrow time window after the stimulus presentation.

**Figure 3.**
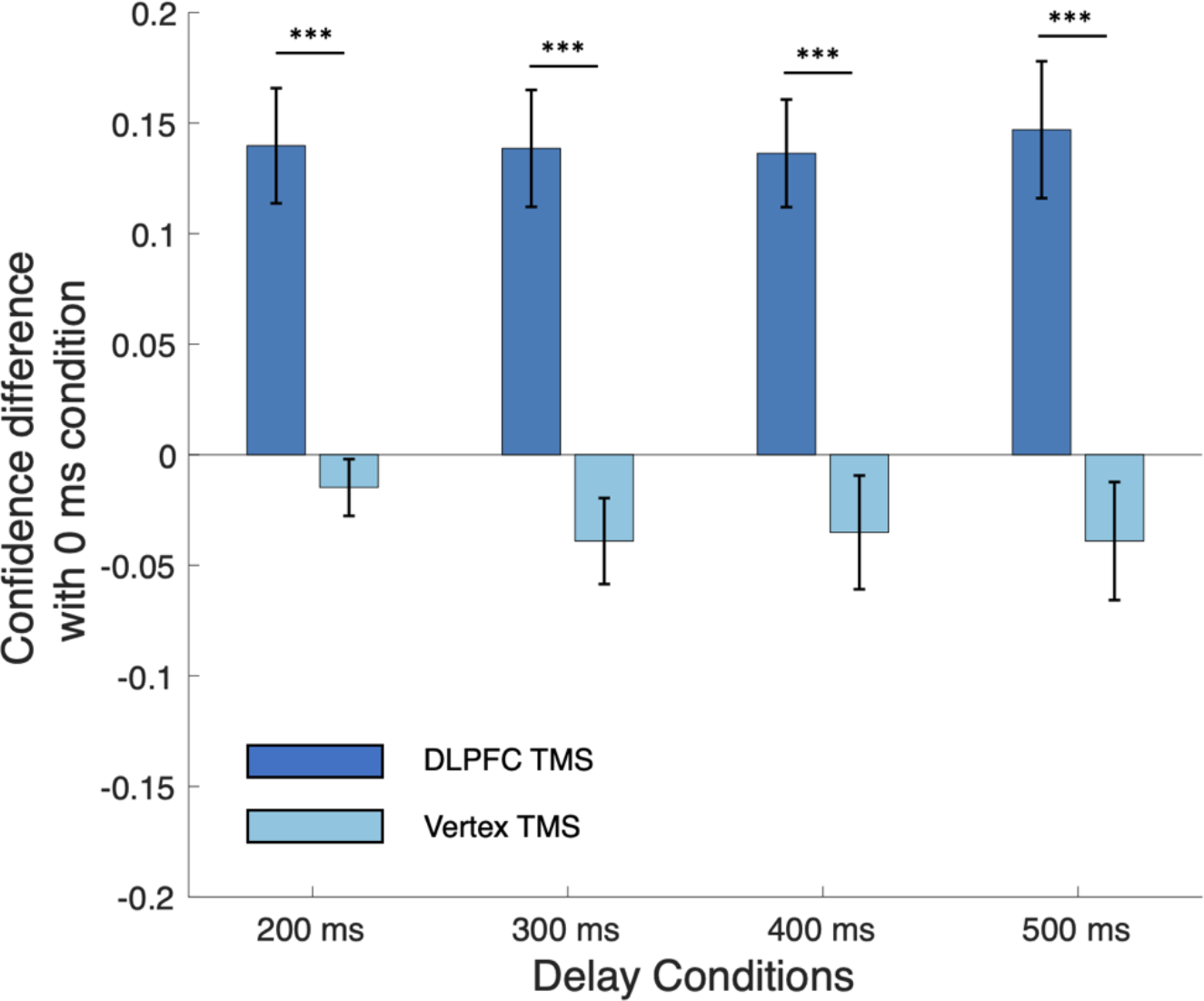
TMS effects on confidence. TMS to DLPFC increased confidence in each delay condition compared to the 0-ms baseline condition, whereas TMS to vertex did not affect confidence for any delay condition compared to the 0-ms baseline. Critically, the effects for both DLPFC and vertex TMS were equivalent across the four delay conditions. Error bars represent SEM; ***, *p* < .001.

**Figure 4.**
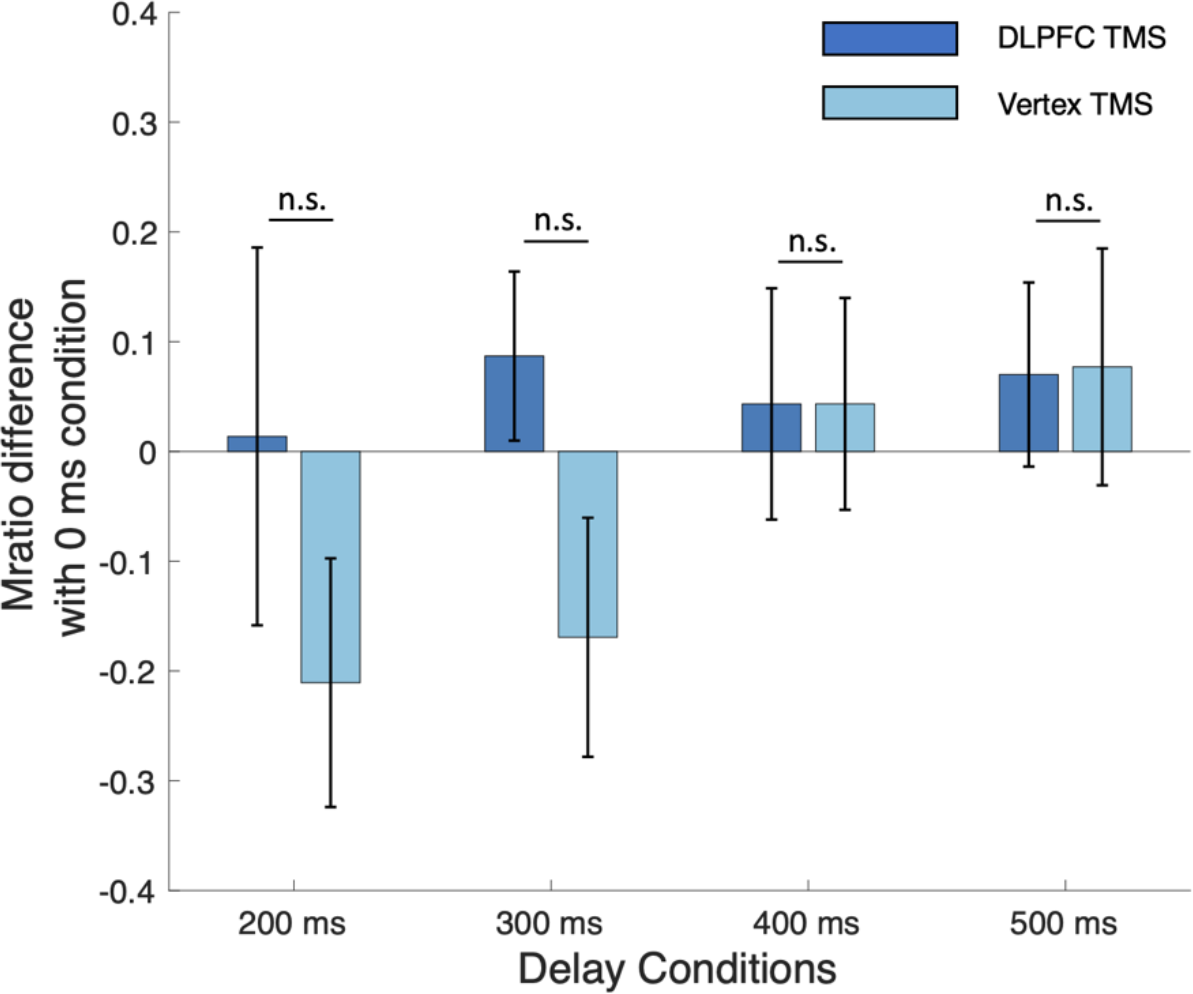
TMS effects on Mratio. TMS to both DLPFC and vertex did not affect Mratio values when compared to the 0-ms baseline condition. There was also no significant effect of TMS site, delay condition, or the interaction between TMS site and delay condition. Error bars represent SEM; n.s., *p* > .05.

Finally, we examined whether TMS affected metacognitive efficiency Mratio. We performed a two-way repeated-measures ANOVA on Mratio difference between the delay conditions and the 0-ms condition with TMS site (DLFPC vs. vertex) and delay condition (200, 300, 400, 500 ms) as factors. We found no main effect of TMS site (F_(1,180)_ = 1.75, *p* = 0.19), no main effect of delay condition (F_(1,180)_ = 0.77, *p* = 0.51), and no interaction between delay condition and TMS site (F_(3,180)_ = 0.62, *p* = 0.60). Pairwise comparisons between the DLPFC TMS and vertex TMS groups for each delay condition also showed no differences in Mratio difference scores (all four *p*’s > 0.06). These results demonstrate that, in line with previous findings (Shekhar & Rahnev, 2018), online TMS to DLPFC has no effect on metacognitive ability.

## Discussion

Understanding the timing of the confidence computations is critical to uncovering the underlying mechanisms of human metacognition. However, despite much progress in other aspects of metacognitive judgments, exactly when confidence is computed is still unclear. To address this question, we tested whether single-pulse TMS delivered to DLPFC between 200 and 500 ms after the stimulus onset would affect either confidence. We found that TMS to all four delay conditions significantly increased confidence in the DLPFC group, but not in a control group where TMS was delivered to the vertex. Importantly, there was no difference in the level of confidence increase among the different delay conditions. Our results demonstrate that confidence is computed over a relatively wide time interval that begins as early as 200 ms after stimulus onset.

Our findings provide evidence against strong versions of post-decisional models of confidence where all confidence-related computation is assumed to take place after a decision has already been made. Several such models may fall into this category. For example, the 2-stage dynamic signal detection (2DSD) model postulates an initial accumulation-to-bound stage that determines the decision, and a second confidence accumulation stage that determines the confidence rating (Pleskac & Busemeyer, 2010). Similarly, the collapsing confidence boundary (CCB) (Moran et al., 2015) and the recent model developed by Herregods et al. (2023) also postulate a similar 2-stage process where no confidence information appears to be computed before the initial decision has been made (Herregods et al., 2023). In the current study, we found that TMS delivered as little as 200 ms after stimulus onset can change the confidence rating without affecting the stimulus sensitivity. Given that average RT was over one second, the internal decision is likely to have been made in less than 200 ms on only a very small percentage of trials (if any). Therefore, our results are incompatible with models that assume that no confidence-related process occur before the decision has been made. Importantly, many models of confidence using the accumulation-to-bound framework assume that confidence signals are present during the initial process of accumulation (Dotan et al., 2018; Hellmann et al., 2022; Rahnev et al., 2016; Ratcliff & Starns, 2009, 2013; Vickers, 1979; Yu et al., 2015) and even the models discussed above could potentially be made compatible with our results by postulating confidence-related processes that begin before the decision has been made.

To be clear, our results do not question the existence of post-decisional processes that contribute to the confidence rating. There is ample empirical evidence that information presented after the decision has been made influences the resulting confidence rating, especially when confidence is given after the initial decision (Desender, Ridderinkhof, et al., 2021). Such findings parallel other literature that post-decisional evidence can lead to changes in the decision itself too (Resulaj et al., 2009). Our findings are perfectly consistent with the existence of post-decisional influences on confidence, but they appear at odds with the idea that confidence is exclusively computed on signals arriving after the decision has been made.

Our findings are most consistent with theories that postulate that confidence is continuously computed in an online fashion during the initial stage of evidence accumulation (Dotan et al., 2018). For example, Dotan et al. (2018) employed a task where subjects continuously indicated their evolving decision using their finger and found that different finger kinematics (position vs. speed) reflected momentary decision and confidence variables independently of each other. A prolonged process of confidence evaluation that roughly overlaps with the decision process fits well with our findings that TMS delivered between 200 and 500 ms after stimulus onset has comparable effects on confidence judgments.

It should be noted that our finding that single-pulse DLPFC TMS delivered after the stimulus onset increased confidence goes in the opposite direction of the results of our previous study where a train of three pulses delivered to DLPFC decreased confidence (Shekhar & Rahnev, 2018). One possible explanation for these different results is that the single TMS pulse in the current study led to excitation that resulted in higher confidence ratings, whereas the TMS train in the Shekhar and Rahnev study led to inhibition that resulted in lower confidence ratings (Caparelli et al., 2012; Romero et al., 2019). It is indeed well known that different TMS parameters can lead to opposite behavioral effects (Caparelli et al., 2012; Huang et al., 2005; Klomjai et al., 2015). Regardless of the underlying reason for these two studies finding effects in the opposite direction, both studies support the notion that DLPFC is a critical node for confidence computation.

In conclusion, we found that single-pulse TMS to DLPFC delivered between 200 and 500 ms after stimulus onset increases confidence, but that a similar effect does not occur for vertex TMS. These results suggest that confidence computation takes place during a broad time window

## Supporting information

Supplementary results and figures

## Acknowledgements

This work was supported by the National Institute of Health (award: R01MH119189) and the Office of Naval Research (award: N00014-20-1-2622).

