## Supplementary results and figures for "The timing of confidence computations in human prefrontal cortex"

**Supplementary Results**

Here we report the results of our preregistered analyses. As per our preregistration, for all of the analyses, we first divided the dataset into an exploratory set (about 1/3 of the data) and a confirmatory set (about 2/3 of the data). In specific, we categorized trials 2, 4, 6 in each group of eight trials to be part of the exploratory set, and included the rest of the trials (1, 3, 5, 7, and 8) in the confirmatory set. Then, we used the exploratory set to develop the hypothesis that we tested using the confirmatory set. We tested four hypotheses defined in our preregistration. Note that we expected TMS to mostly decrease confidence, and hence preregistered the analyses with that expectation, but below we report separately analyses aimed at both increased and decreased confidence.

First, we examined whether TMS delivered several hundred ms after stimulus presentation would lead to a significant change in confidence compared to the baseline *for at least a subset of subjects*. Namely, we first computed the average confidence for the 0-ms condition, as well as for the four TMS delay conditions (200, 300, 400, 500 ms) in the exploratory dataset. For each subject, we then identified the delay condition with highest average confidence. Subjects for whom the average confidence in the delay condition was at least 0.1 higher than the average confidence in the 0-ms condition were passed to the second stage. In the second stage, we analyzed that data in the confirmatory set and examined whether the confidence in the 0-ms condition was lower than the selected delay condition across all passed subjects. A total of 37 subjects were selected based on the passing criterion. A one-sided paired t-test showed that for the selected subjects and timepoints from the exploratory set, there was a significant increase in confidence in the confirmatory set too (t(36) = 6.36, *p* = 1.15 X 10^-7^). We repeated the same procedure to test for consistent confidence decrease. Our procedure, however, only resulted in selecting two subjects in the exploratory set, and a one-sided paired t-test showed no significant decrease in the confirmatory set (t(1) = 3.33, *p* = 0.91). We repeated the same analyses for Mratio and passed 34 subjects to test for Mratio increase. A one-sided paired t-test showed that for these 34 selected subjects and timepoints from the exploratory set, there was a small but significant increase in Mratio in the confirmatory set (t(33) = 1.75, *p* = 0.04). When we repeated the procedure to test for Mratio decrease, nine subjects were passed but we didn’t find a significant decrease in Mratio in the confirmatory set (t(8) = -0.08, *p* = 0.47).

Second, we tested whether TMS delivered several hundred ms after stimulus presentation would lead to a significant increase or decrease in confidence compared to the 0-ms condition *for the group as a whole*. Namely, we followed the same procedure but tested all subjects in the second stage. In other words, in the first stage, for each subject, we selected the delay with maximum confidence using the exploratory set. Then, in the second stage, we used a one-sided paired t-test to test whether the baseline confidence was lower than the selected delay condition in all subjects. We found a significant increase in confidence for the group as a whole in the confirmatory set (t(42) = 6.8, *p* = 1.37 X 10^-8^). However, a similar analysis on confidence decrease revealed no significant effect (t(42) = 7.40, *p* = 1). We repeated these analyses for Mratio, and we found that neither the increase nor the decrease were significant (for increase: t(42) = 0.79, *p* = 0.22; for decrease: t(42) = 1.73, *p* = 0.95).

Third, we tested whether TMS delivered at a specific time point after stimulus presentation would lead to a significant increase or decrease in confidence compared to the 0-ms condition for the *group as a whole*. Specifically, we followed the same procedure as for Hypothesis 2 but selected the same delay condition for all subjects. The delay condition was selected as the one with the lowest or highest average confidence in the exploratory set. Again, a one-sided paired t-test was used to compare that delay condition to the 0-ms condition in the confirmatory set. We found that the 300-ms condition had the highest confidence in the exploratory set (0 ms: 2.31; 200 ms: 2.47; 300 ms: 2.49; 400 ms: 2.48; 500 ms: 2.49). Testing the 300-ms condition in the confirmatory set, we found that it featured significantly higher confidence than that the 0-ms condition (t(42) = 5.96, *p* = 2.3 X 10^-7^). On the other hand, the 200-ms condition had the lowest confidence rating among the delay conditions in the exploratory set, but it did not show a significant increase in the confirmatory set (t(42)= 6.06, *p* = 1). For Mratio, we identified the 500-ms condition as the condition with the maximum Mratio, and the 200-ms condition as the one with the minimum Mratio in the exploratory set. However, one-sided t-test showed that neither of them was significant in the confirmatory set (testing for an Mratio increase in the 500-ms condition: t(42) = 1.12, *p* = 0.13; testing for an Mratio decrease in the 200-ms condition: t(42) = 0.93, *p* = 0.82).

Fourth, we tested whether the effect of TMS on confidence would differ across the delay conditions of 200 to 500 ms. To examine this, we performed a repeated-measure ANOVA on the 4 delay conditions. In our preregistration, we stated that if the ANOVA was significant (demonstrating that there were significant differences between the different delay conditions), then we would follow up with two-sided paired t-tests to explore which conditions differed significantly from each other. Note that the procedure of this analysis is the same as the analysis we reported in the main paper, but the results we are reporting here are results without any exclusion based on accuracy. As in the main paper, we found that there was no difference in confidence across the four delay conditions (F_(3,126)_ = 0.67, *p* = 0.57). Similar results were obtained for Mratio as well (F_(3,126)_ = 0.60, *p* = 0.62). Because the ANOVA was not significant, we did not follow up with any paired t-tests.

Overall, the results of these preregistered analyses are consistent with the results we report in the main paper. Specifically, both sets of analyses point to the conclusion that TMS to DLPFC resulted in a confidence increase across all four delay conditions, and that there was no difference in the confidence increase among these four delay conditions.

**Supplementary Figures**

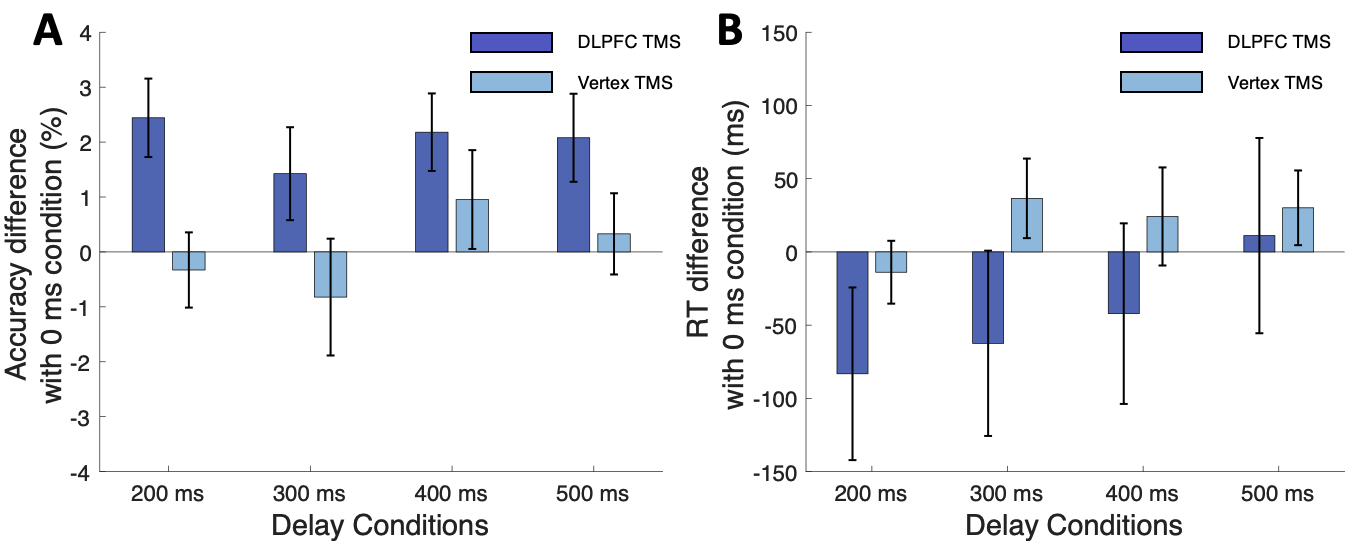

**Supplementary Figure 1. TMS effects on accuracy and RT without excluding subjects with low accuracy in the 0-ms condition.** (A) The effect of TMS on the accuracy difference between each delay condition and the 0-ms condition. A two-way repeated measures ANOVA on accuracy difference between each delay condition and the 0-ms condition with factors TMS site (DLPFC and vertex) and delay conditions (200, 300, 400, and 500 ms) showed a main effect of TMS site (F_(1,240)_ = 9.56, *p* = 0.0022), but no main effect of delay condition (F_(3,240)_ = 0.68, *p* = 0.57), and no interaction between TMS site and delay condition (F_(3,180)_ = 0.26, *p* = 0.85). These results are in line with our findings in the main analyses with the exception of the higher accuracy differences for the DLPFC condition that are a direct effect of not excluding the subjects with low accuracy in the 0-ms condition. (B) The effect of TMS on RT difference between each delay condition and the 0-ms condition. A similar two-way repeated-measures ANOVA on the RT difference between each delay condition and the 0-ms baseline condition showed no effect of TMS site (F_(1,240)_ = 1.73, *p* = 0.19), delay condition (F_(3,240)_ = 0.35, *p* = 0.79), or an interaction between the two (F_(3,240)_ = 0.12, *p* = 0.95). Together, these results confirm that there were no differences among the four delay conditions in either accuracy or RT regardless of whether subjects with low accuracy in the 0-ms condition are excluded or not. Error bars represent SEM.

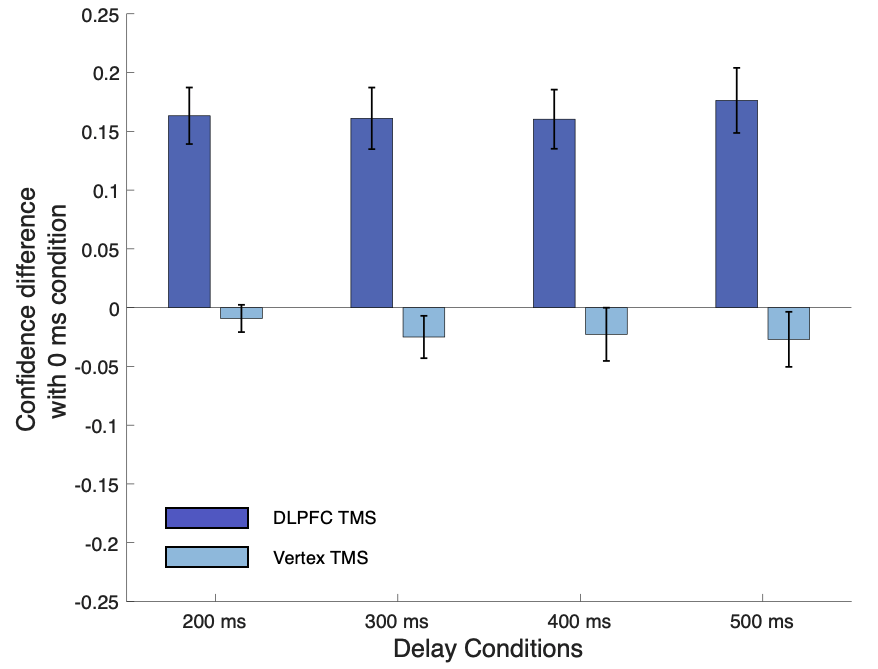

**Supplementary Figure 2. TMS effects on confidence without excluding subjects with low accuracy in the 0-ms condition**. TMS to DLPFC increased confidence in each delay condition compared to the 0-ms baseline condition (all *p*’s < 0.001), whereas TMS to vertex did not affect confidence for any delay condition compared to the 0-ms baseline (all *p*’s > 0.18). A two-way repeated-measures ANOVAs on the confidence difference between each delay condition and the 0-ms condition with factors TMS site (DLPFC and vertex) and delay conditions (200, 300, 400, and 500 ms) showed that there was a significant main effect of TMS site on confidence (F(1,240) = 82.23, *p* < 0.001), no significant effect of delay condition (F(3, 240) = 0.05, *p* = 0.99), and no interaction between the two (F(3, 240) = 0.1, *p* = 0.96). These results replicate the results in our main analyses and again confirm that there were no differences among the four delay conditions regardless of whether subjects with low accuracy in the 0-ms condition are excluded or not. Error bars represent SEM.

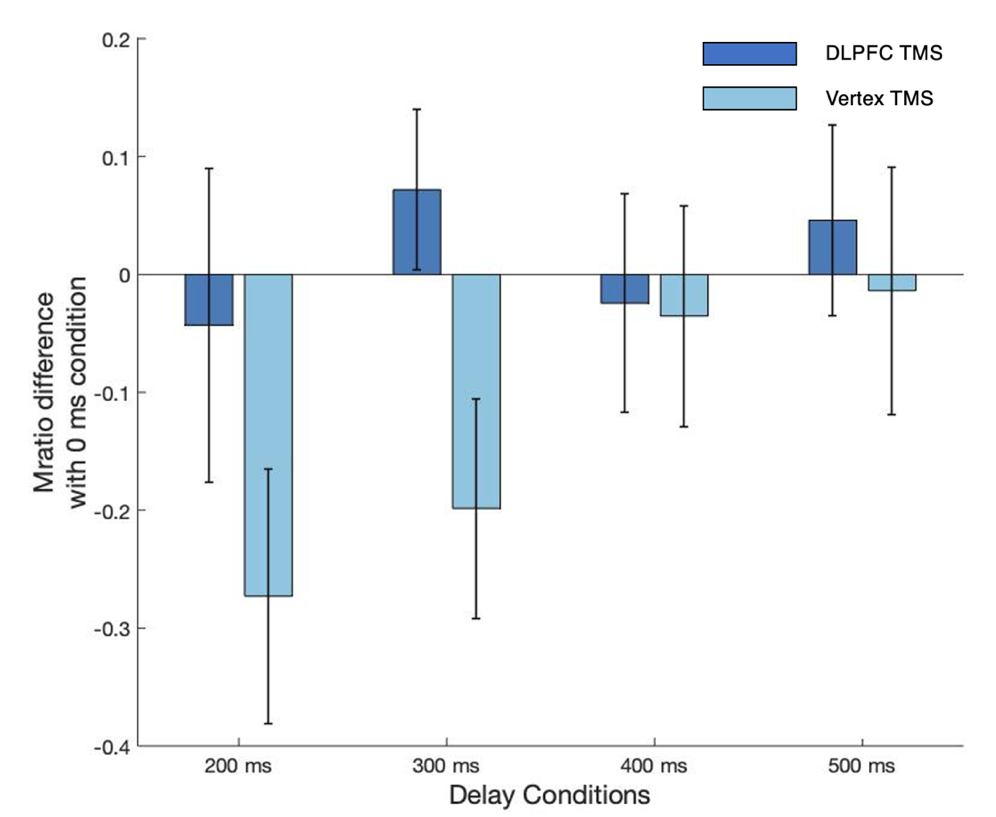

**Supplementary Figure 3. TMS effects on Mratio without excluding subjects with low accuracy in the 0-ms condition.** A two-way repeated-measures ANOVAs on the Mratio difference between each delay condition and the 0-ms condition with factors TMS site (DLPFC and vertex) and delay conditions (200, 300, 400, and 500 ms) showed that there was a significant main effect of TMS site on Mratio (F(1, 240) = 3.17, *p* = 0.08), no significant effect of delay condition (F(3, 240) = 0.85, *p* = 0.47), and no interaction between the two (F(3,240)=0.63, *p* = 0.60). These results confirm that there were no differences among the four delay conditions regardless of whether subjects with low accuracy in the 0-ms condition are excluded or not. Error bars represent SEM.
